# MitoMouse is a model reconstruction of murine mitochondrial metabolism

**DOI:** 10.1101/2023.01.02.522468

**Authors:** Stephen P. Chapman, Thibaut Molinié, Arnaud Mourier, Bianca H. Habermann

**Author notes:** corresponding authors : Stephen P. Chapman, Bianca H. Habermann.

## Abstract

Mitochondria, apart from being the powerhouses of our cells, are key players in cellular metabolism and homeostasis. As a consequence, the core bioenergetics and metabolism of mitochondria are well studied considering the role that dysregulated mitochondrial metabolism plays in disease. Flux Balance Analysis, used in conjunction with metabolic model reconstructions is a powerful computational tool that predicts metabolism. The resulting quantitative descriptions of metabolic dysregulation not only allows current hypotheses to be tested *in silico*, but can also lead to novel model driven hypotheses that can be experimentally verified. Several metabolic reconstructions for human and mouse metabolism exist, but no model specific to mouse mitochondrial metabolism exists. Here, we have created a mouse-specific mitochondrial metabolic model, mitoMouse, which is based on the high-quality human MitoCore mode. MitoMouse contains 390 genes and 445 metabolites involved in 560 unique reactions, is able to model central carbon metabolism and has been extended to contain reduction of the CoQ complex of Oxidative Phosphorylation by the enzyme DHODH. MitoMOuse was validated to accurately model the important metabolic switch involving CoQ reduction resulting from increased malate import, as recently shown in mouse cardiac tissue. We expect this model to be of immense interest and relevance to researchers working on murine mitochondrial metabolism.

## Introduction

Mitochondria are essential organelles present in nearly all eukaryotic cells. One of their main roles is to provide cellular energy in the form of ATP through oxidative phosphorylation (OXPHOS). OXPHOS involves the transfer of electrons from a number of electron donors, along the respiratory chain (RC), historically described as composed of four enzyme complexes (complexes I–IV). The traffic of electrons through the RC is mediated by the mobile electron carriers, Coenzyme Q (CoQ) and cytochrome C. Electron transfer through each complex is coupled to proton translocation from the mitochondrial matrix to the intermembrane space which generates a proton motive force (PMF) across the inner membrane that is used by ATP-synthase, to phosphorylate ADP to ATP ^1^. Several other reduced metabolites are also directly oxidised by the RC through oxidoreductases transferring electrons to CoQ. In mammals, these are mitochondrial glycerol-3-phosphate dehydrogenase (GPD2/mG3Pdh), dihydroorotate dehydrogenase (DHODH) ^2^ and the electron transfer flavoprotein (ETF-Qo) dehydrogenase, the first step of the mitochondrial fatty acid ß-oxidation. The presence of these oxidoreductases positions the electron carrier CoQ at the crossroad of several key metabolic pathways (Krebs cycle, pyrimidine synthesis and fatty acid ß-oxidation).

In addition to producing ATP, OXPHOS is metabolically essential to regenerate the oxidised redox cofactor NAD□ from NADH, which along with ATP, are indispensable cofactors for all metabolic pathways. The malate aspartate shuttle (MAS) is the main shuttle that traffics cytosolic NADH (produced from central carbon metabolism) into mitochondria and contributes to mitochondrial maintenance of NADH/NAD^+^ redox cofactor balance. The MAS has been shown to be the predominant NADH redox shuttle in highly oxidative tissues such as the brain, heart and liver ^3^. This key redox shuttle involves mitochondrial carriers as well as cytosolic and mitochondrial malate dehydrogenases (MDH1, MDH2) and glutamate aspartate transaminases (GOT1, GOT2) ^4^. Intercompartment metabolite cycling through the MAS allows MDH1 and MDH2 to work in opposite directions (oxidising cytosolic NADH and reducing mitochondrial NAD^+^). This enables CI to indirectly oxidise cytosolic NADH. Beyond NADH redox maintenance through CI, the RC is also key to sustaining the tricarboxylic acid cycle (TCA cycle) via CII. Finally, mitochondria are also responsible for the production of reactive oxygen species (ROS), a by-product from OXPHOS, which also serves as an essential secondary messenger ^5^.

Defective mitochondria and dysregulated mitochondrial metabolism, either due to mutation of mitochondrial or nuclear DNA, or due to other factors, are known to play a crucial role in many disorders. Beyond these disorders, altered mitochondrial metabolism also contributes to many other diseases, such as neurological diseases, or cancer ^6,7^ and plays a crucial part in ageing^8,9^. Studying mitochondrial metabolism and the consequences of its perturbation is therefore important to identify the key players involved in the onset or progression of many different human disorders, as well as ageing.

The mitochondrial genome, metabolism and function are highly conserved between human and mouse^10,11^. The genetic and physiological similarity between mice and humans make mice therefore suitable animal models to study mitochondrial structure, function and regulation, as well as human (mitochondrial) diseases, leading to potential novel therapeutic strategies. Although there are considerable differences between mice and humans, particularly regarding genetics, physiology, and immunology^12^. Laboratory mouse models are unavoidable to investigate complex pathogenic mechanisms and evaluate therapeutics strategies. The combination of multiple factors such as cost, availability, translatability of the results, ease of manipulation, and ethical implications, rationalise the fact that laboratory mouse is the most widely used mammalian animal in biomedical research. The International Mouse Phenotyping Consortium has generated, phenotyped and archived more than 6000 knockout mice on the C57BL/6 background, the most well-known and widely used inbred mouse strain^13^. Beyond being a valuable model to study human disorders, laboratory mouse models are also highly relevant to elucidate tissue specific changes adapting energy metabolism to tissue specific needs. Like cell morphology, cell energy metabolism of the differentiated cell types is highly remodelled to fit their functional specialisation.

The recent discovery of an evolutionary conserved mechanism orchestrating the RC electron fuelling at the CoQ-oxidoreductase crossroad shed new light on the metabolic role played by mitochondria^14–16^. In mouse mitochondria, allosteric regulation by MAS produced malate, and subsequent conversion to oxaloacetate (OAA) reduced complex II (CII) activity due to increasing complex I (CI) activity, a feature first shown in yeast and thus, conserved in evolution^15–19^. As a consequence, electron flow from succinate towards NADH oxidation was rewired. This was triggered by physiological concentration of malate. These results suggest that oxidation of imported malate by MDH2 rewires the fuelling of the RC from CII to CI, facilitated by CoQ in its role as a common substrate to both complexes^16^. The discovery that malate import via MDH1 can modulate the activity of CoQ constitutes a major conceptual breakthrough in the field of mitochondrial energy metabolism: this novel regulatory ‘CoQ-contest’ mechanism enables the RC to prioritise metabolic flows by orchestrating the activities of coenzymes Q oxidoreductases.

Interestingly, independent work demonstrated that the CoQ metabolic crossroad is heavily reshuffled across tissues through important tissue-specific variations in CoQ levels^20^, and through stringent tissue-specific expression of the different CoQ-oxidoreductases^21^. This suggests that the CoQ-contest could be key to rewire RC electron fuelling in a tissue-specific manner. The recent observation that CoQ-oxidoreductases exhibit diverging abilities to form stable structural assemblies with other RC complexes puts a new focus on supramolecular organisation and its potential role in orchestrating electron fuelling at the CoQ crossroad of the RC^22^. Yet, the functional relevance of the RC supramolecular organisation and its role in controlling the RC electron fuelling is still under debate^23–26^.

The genomes of many organisms and organelles have already been sequenced and gene expression profiles are constantly being determined in many conditions and model systems^27^. Mass spectrometry-based protein surveys have quantified the global proteome^28^ and metabolome^29^ on cellular and systems levels. This enormous amount of information at different scales of organisation can be used to obtain a new perspective that connects a genotype to a phenotype, through metabolism^30^, by means of genome-scale models (GSM’s) that contain all known metabolic reactions encoded by the system’s metabolic genes. Each enzyme-associated reaction within the GSM is encoded by a gene-protein-reaction (GPR) rule describing the relationship between all necessary genes to ensure production of a gene product involved in metabolism. The encoded metabolic pathways can be constructed using our knowledge of metabolic genes, to produce metabolic models at genomic scale. Flux Balance Analysis (FBA) can then be used to simulate a GSM and predict metabolic fluxes that can determine an observed phenotype. Since the advent of the first GSM that described *H. influenza* metabolism^31^ over 6000 metabolic reconstructions have been produced, covering a wide range of species from bacteria, and archaea to complex eukaryotes^32^.

The mitochondrial metabolism of human and mouse is included in several genome-scale reconstructions, for instance the Recon 1 model, which was the first generic human GSM published^33^. An orthologous mouse model (iMM1415) was then created from the human model^34^. Recon 1 was updated to Recon R2, which included additional biological information and the correction of various modelling errors such as Recon 1’s inability to correctly predict realistic ATP yields^35^. Recon R2 was subsequently upgraded to Recon 3D, which includes more reactions, metabolites and genes and extensive human GPR associations^36^. From this model, an updated mouse model, iMM1865 was produced using a top-down orthology-based methodology by mapping human genes of Recon 3D to mouse^37^.

One challenge facing predictive modelling at the genome-scale level lies in the fact that large models encompassing thousands of reactions are more prone to error than smaller and more concise models. This is a consequence of missing knowledge and annotation and sources of error include incorrect parameterisation of reaction directionality constraints and issues relating to the incorrect compartmentalisation of reactions and metabolites. Using genome-scale models to simulate complex diseases can therefore result in erroneous circuits and thus lead to mispredictions. Furthermore, all existing models of human, or mouse global metabolism neglect to predict the proton motive force (PMF) that drives the phosphorylation of ADP to ATP within mitochondria. This limits their use in accurately modelling mitochondrial metabolism and bioenergetics. Several models of human mitochondrial metabolism exist^38,39^, with MitoCore representing the latest and most comprehensive model of human cardiomyocyte mitochondrial metabolism^38^. MitoCore includes the RC within its reconstruction and is able to accurately model the PMF and subsequent ATP production, as well as a more accurate partitioning of reactions between the cytosol and mitochondria. This model has successfully been applied to model fumarase deficiency^38^, impaired citrate import^40^ and predicted an accurate respiratory quotient following metabolism on glucose and palmitate substrates^41^. This demonstrates MitoCore’s ability to accurately model human cardiomyocyte mitochondrial metabolism.

In this work, we set out to construct a mouse model of mitochondrial metabolism, based on the human MitoCore model. We first converted the human MitoCore to mitoMouse, a mouse-specific model of mitochondrial metabolism, to allow the modelling of mitochondrial metabolism in this organism using FBA. We also expanded the model to include CoQ reduction from DHODH, introducing the importance of CoQ and the RC in orchestrating metabolic paths in mitochondria. We tested our model on experimental data provided by ^16^, who showed that increased import of malate through the Malate Aspartate Shuttle (MAS) leads to increased fluxes associated with oxaloacetate metabolism and results in the prioritisation of Complex I reduction over Complex II. MitoMouse accurately predicts mitochondrial ATP production following glucose oxidation, but also the switch in CoQ reduction from CII to CI following malate import and metabolism into OAA. We updated mitoMouse to conform with the latest SBML level 3 (v1) using Python (v3) and provide a tutorial in the form of a freely available Jupyter notebook (https://gitlab.com/habermann_lab/mitomouse).

## Methods

### Mapping MitoCore genes to mouse orthologs

Considering the similarity between human and mouse mitochondria, conversion of MitoCore’s gene-product-reaction (GPRs) to mouse equivalents would enable the construction of a mouse-specific model of mitochondrial metabolism. Human genes contained within MitoCore were mapped to corresponding mouse orthologs by automatic application programming interface (API) programming complemented with a manual search of Biomart^42^ and the ENSEMBL database, as well as with orthology information stored within MitoXplorer2^43^. This resulted in 388 mouse orthologs out of the original complement of 391 MitoCore genes. Figure 1 a summarises the construction of mitoMouse from the MitoCore model. The set of mitoCore GPR rules were compiled to their corresponding logical expressions for mitoMouse based on orthology relations between human and mouse genes (Supplementary Table S1 b).

**Figure 1:**
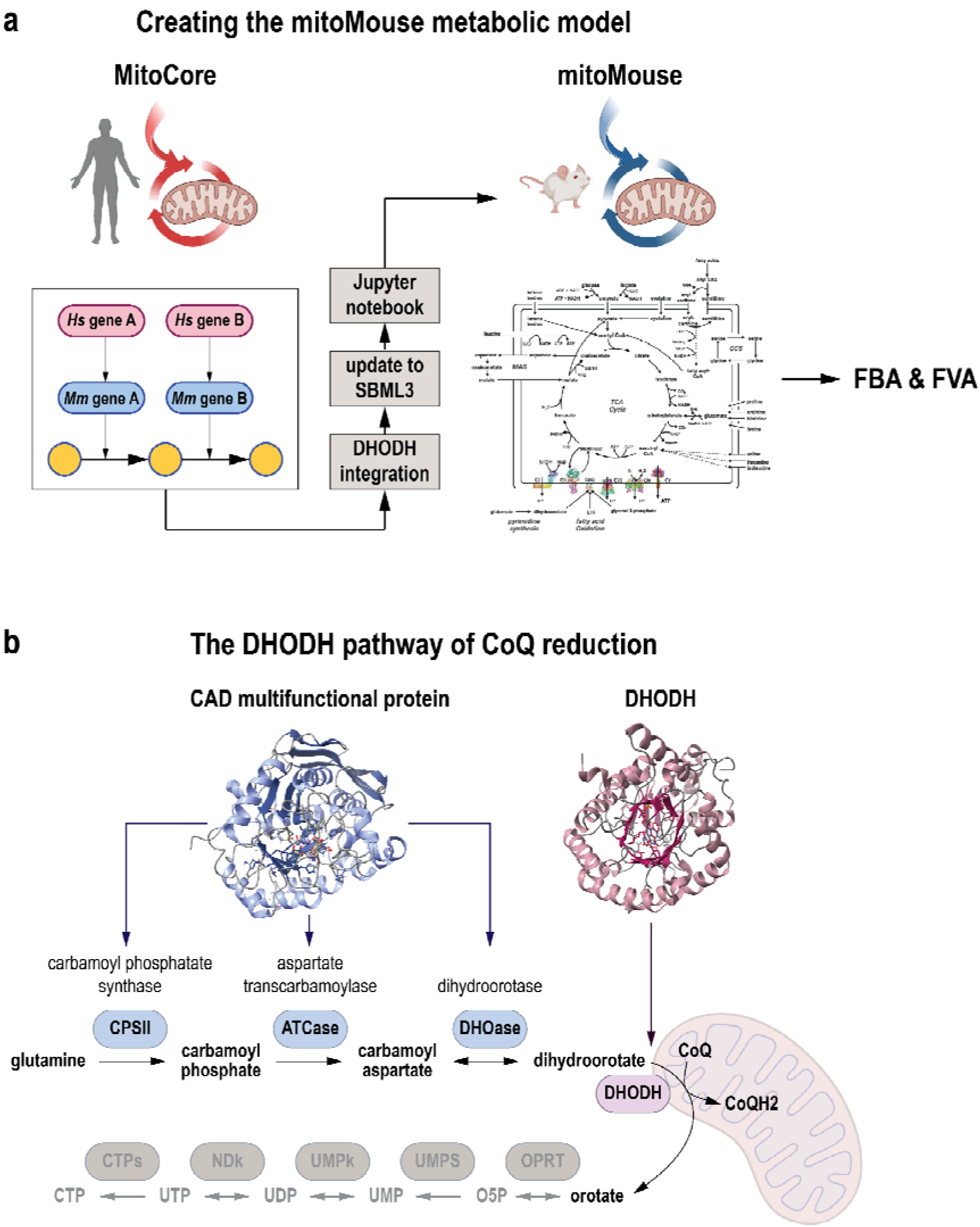
**(a)** MitoMouse construction process. The MitoCore model of human cardiac mitochondrial metabolism was used as the base model for mitoMouse. All human genes were mapped to mouse orthologs and the missing pathway of CoQ reduction by DHODH was integrated into the model. MitoMouse was encoded to SMBL level 3 (version 1) standards and extended using the Flux Balance Constraints package. It provides the standardised format for the encoding, exchange and annotation of constraint-based models. **(b)** DHODH expansion of mitoCore involves addition of 5 new reactions encoded by 2 genes and 4 new metabolites.

### DHODH expansion

MitoCore was missing DHODH reduction of CoQ, yet contained glutamine metabolism which is the starting substrate for *de novo* pyrimidine synthesis. This metabolic pathway involves the initial conversion of glutamine to carbamoyl phosphate facilitated by carbamoyl phosphate synthase. Carbamoyl phosphate is then metabolised to carbamoyl aspartate through the activity of aspartate carbamoyltransferase, which is subsequently metabolised into dihydroorotate by the enzyme dihydroorotase. These three enzymes are collectively part of a single multifunctional protein abbreviated as CAD (Carbamoyl Aspartate Dihydroorotase). Dihydroorotate then reduces CoQ to produce reduced CoQ and orotate, facilitated by the enzyme dihydroorotic acid dehydrogenase (DHODH) that sits at the surface of the outer mitochondrial membrane. As such, orotate is never imported into the mitochondria and remains cytoplasmic. We included these metabolic reactions and new metabolites in mitoMouse (Figure 1 b). Orotate removal from the model was implemented by the addition of a demand reaction to maintain flux consistency. In total, five new reactions were added that incorporate four new metabolites and two new genes.

### Update to SBML3 (version 1)

The original MitoCore model was encoded using SBML level 2 annotation, which is now considered outdated. We updated the mitoMouse to the most recent, relevant specification of SBML level 3 (version 1)^44^ (Supplementary File S1).

### Constraint-based modelling using FBA and FVA

Flux Balance Analysis (FBA) and Flux Variability Analysis (FVA) were performed using Python in conjunction with the COBRApy toolbox^45^, using the default ‘GLPK’ solver. mitoMouse was first tested on its ability to correctly produce accurate ATP levels from glucose oxidation. All nutrient input reactions except glucose and oxygen were constrained to zero to reflect aerobic glycolytic conditions. Maximisation of ATP production was used as the objective function.

A robustness analysis was performed to investigate the ability of mitoMouse to import malate through the Malate-Aspartate Shuttle (MAS) and subsequent metabolism into oxaloacetate. The constraints for the malate import reaction from MAS varied from 0 to a maximum of 3 mmol/gDw/h at 0.1 step intervals. The resulting flux through complex I and complex II reactions were then observed as a function of imported malate. The same constraints were applied as previously described and the ATP-producing reaction was optimised.

The resulting flux distributions for malate import were initially mapped using the Escher package^46^, redrawn for clarity, and compared (Jupyter notebooks containing constraints, simulation parameters and Escher flux visualisations are provided in Supplementary File 2). The resulting fluxes from the two conditions were scored and ranked in decreasing order, based on their absolute difference (Supplementary Table S1 c).

### Reaction Essentiality Analysis using Flux Variability Analysis

FVA is an FBA-based method for characterising all feasible states of genome-scale metabolic models that satisfy a given objective function^47^. Numerous combinations of flux vectors can satisfy the given objective function by using different pathways, resulting in several potential flux distributions. As such, the points in the solution space that maximise or minimise a given objective function can be characterised^48^. FVA was used to compute minimal and maximal possible fluxes obtained for each reaction within mitoMouse, whilst constraining the ATP-producing reaction to the rate obtained from FBA (flux values from FVA are contained in Supplementary Table S1c). For each reaction, the minimal possible flux under 3 mmol malate import (*a*) was subtracted from maximal fluxes under zero malate import (*b*). Conversely, minimal fluxes under zero malate import conditions (*c*) were subtracted from maximal fluxes of malate imported at 3 mmol (*d*).

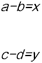

Absolute differences from the above two subtractions were calculated to give the smallest possible difference between conditions for a given reaction to satisfy the objective function of maximising ATP production in each case.

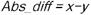

For reactions where there was a clear separation between respective fluxes of zero and maximum malate import, these were classified as non-overlapping fluxes; in other words, there was always a difference in the flux between the two conditions. Meanwhile, overlapping fluxes were given a score of 0 and discarded. All reactions that resulted in positive fluxes were identified and analysed.

To determine which reactions and metabolic pathways are active or dormant in a particular condition, a reaction essentiality analysis was then performed from the non-overlapping reactions derived from FVA. For the two conditions, reactions of mitoMouse were classified as flux essential (requiring a non-zero flux), flux substitutable (where the range of possible fluxes span zero) and flux blocked (reactions with minimum and maximum flux of zero). This method uses FVA results to determine whether a flux is always, potentially or never required for optimal ATP production.

## Results

### The MitoMouse mitochondrial metabolic model

The aim of this work was to produce a mouse-specific metabolic model of mitochondrial metabolism that incorporates new knowledge on CoQ fueling. We first translated the human MitoCore model into mouse using orthology inference, creating the basic mitoMouse model. Key metabolic pathways that include the TCA cycle, the Malate Aspartate Shuttle (MAS); OXPHOS and ATP synthesis; the Glycine Cleavage System (GCS) and fatty acid oxidation are also retained from the original model. MitoMouse now also includes *de novo* pyrimidine synthesis from glutamate leading to the reduction of the CoQ complex by the enzyme DHODH. MitoMouse contains 390 genes and 445 metabolites involved in 560 unique reactions. The complete list of metabolites and reactions are available in Supplementary Table S1a and Supplementary Table S1 b respectively, and summarised in Figure 2 a.

**Figure 2:**
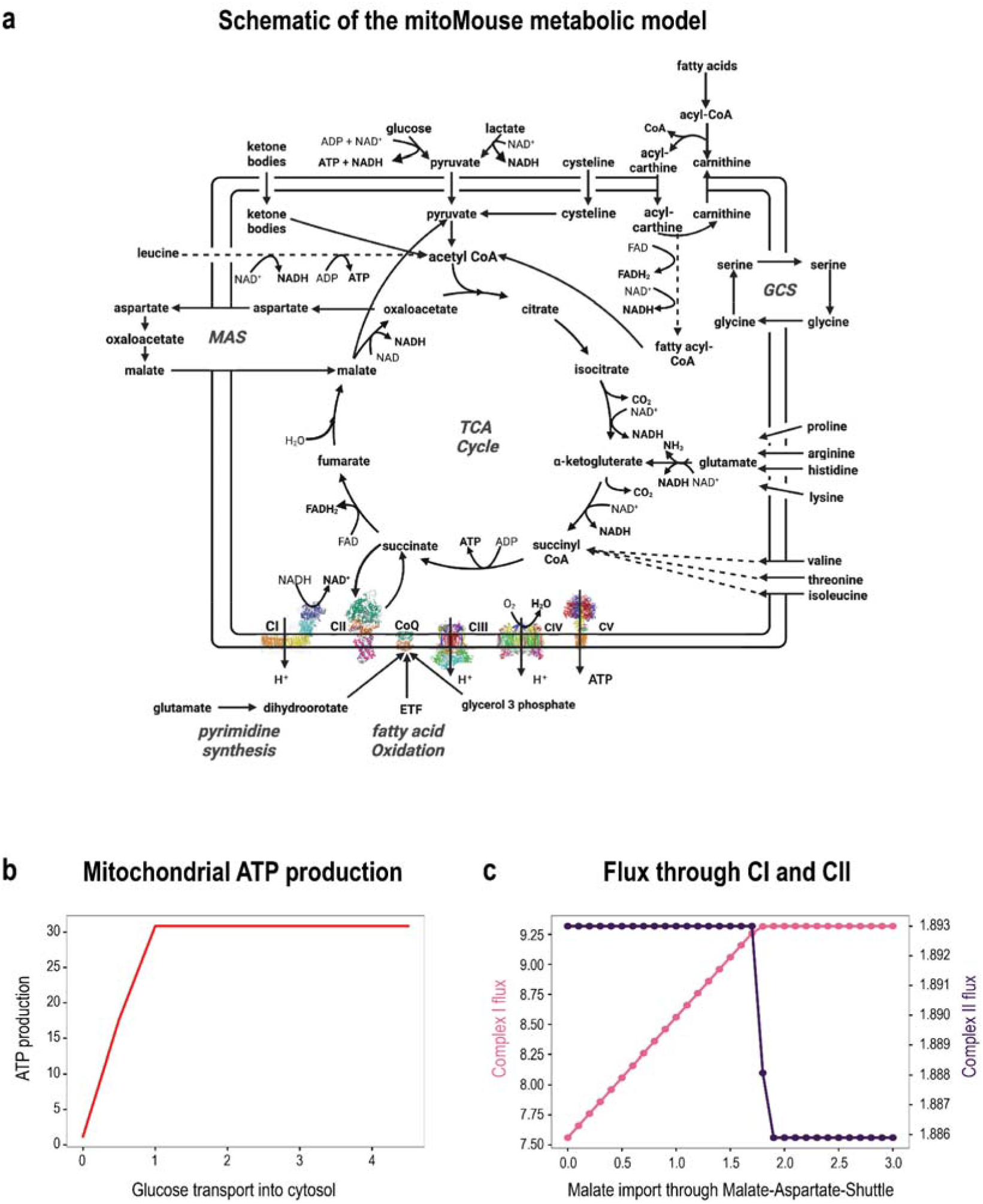
**(a)** Schematic of the mitoMouse model, a complete medium-scale model of murine cardiac mitochondrial metabolism. Abbreviations: CI: Complex 1; CII: Complex 2; CIII: Complex 3; CIV: Complex 4; CV: Complex 5; GCS; Glycine Cleavage System, MAS: Malate Aspartate Shuttle. **(b)** MitoMouse correctly predicts that the metabolism of 1 molecule of glucose produces 31 molecules of ATP. **(c)** Malate import prioritises complex 1 over complex 2. Forcing the import of malate flux through the MAS from 0.0 to 3.0 (mmol/gDw/Hr) predicted a change in preference of electron donor from complex II, to complex I when malate import uptake reached 1.7 (mmol/gDw/Hr).

### Testing mitoMouse to predict known ATP production rates

We next performed Flux Balance Analysis on mitoMouse to investigate the model’s ability to predict accurate ATP levels. As shown in Figure 2b, the model is able to accurately predict ATP yields, with 1 molecule of glucose yielding 31 molecules of ATP. 94% of this ATP is generated from ATP synthase (Complex V) and the remaining 4% of ATP is produced by the TCA cycle enzyme ‘Succinyl-CoA’, (R_SUCOASm) as is expected. This result is well within the reported range of 30 -33 molecules of ATP from the complete oxidation of glucose^49^.

### Predicting CoQ contest using mitoMouse

It was recently observed that malate imported via the Malate-Aspartate Shuttle (MAS) resulted in a switch of electron fueling to CoQ from CII to CI at a physiological concentration of 2mM malate^16^. As another means of model validation, we tested the ability of mitoMouse to capture this switch by increasing the flux of the reaction responsible for malate import through the MAS and observing the flux through CII and CI. At lower malate import fluxes, flux through CII was constantly high and CI flux was increasing with increasing malate concentrations. When malate import reached 1.7 mmol/gDw/Hr, CI flux reached a maximum and flux through CII dropped rapidly. Next to being able to model this switch of electron fueling of CoQ from CII and CI with mitoMouse, the resulting fluxes show a quantitative agreement with the experimental observations that show MAS imported malate triggers a shift from succinate oxidation to NADH oxidation (Figure 2 c).

### Analysis of the flux distribution for low and high malate import

With the validated mitoMouse model, we performed Flux Variance Analysis (FVA) to analyse the flux distributions resulting from FBA and assessed the essentiality status of each reaction associated with the two conditions: 0.0 malate vs 3.0 malate import (mmol/gDw/h) through the malate import (AKGMALtm) reaction.

#### Zero malate import

Constraining the malate import reaction to zero effectively blocked malate import through MAS and resulted in predictions of malate import from alternative sources. The model predicted a rerouting of metabolism by inducing the activity of the compensatory citrate malate shuttle (CMS) to increase malate availability within the TCA cycle at the expense of exporting citrate. Additional malate was also predicted to enter the TCA cycle in exchange for inorganic phosphate by the activity of MALtm, a malate/phosphate antiporter. Following malate entry into the TCA cycle, mitoMouse predicted 99% is converted into OAA through the activity of malate dehydrogenase (MDH2) and 1% of the malate being decarboxylated into pyruvate by the malic enzyme reaction (ME1), followed by increasing pyruvate metabolism into acetyl-CoA and with it, NADPH. Acetyl-CoA was fed into the TCA cycle leading to citrate metabolism which is partly removed by the activity of the CMS, and partly converted to isocitrate by the enzyme aconitase. Isocitrate is metabolised into alpha-ketoglutarate (aKG), producing CO_2_ and reducing power in the form of NADH. A flux of 1.89 mmol/gDw/h was sustained through subsequent TCA cycle enzymes alpha-ketoglutarate dehydrogenase (aKGDH), succinate CoA synthetase (SUCC-CoAS), succinate dehydrogenase (SUCDH) and fumarase (FUM). As a consequence of the flux through aKGDH and SUCDH, NADH and FADH_2_ are produced and made available as substrates that can feed into the reactions of OXPHOS. The model predicted that inhibiting the activity of AKGMALtm to zero reduced the overall flux capacity of the MAS and so low fluxes of aspartate export were observed (Figure 3 a).

**Figure 3:**
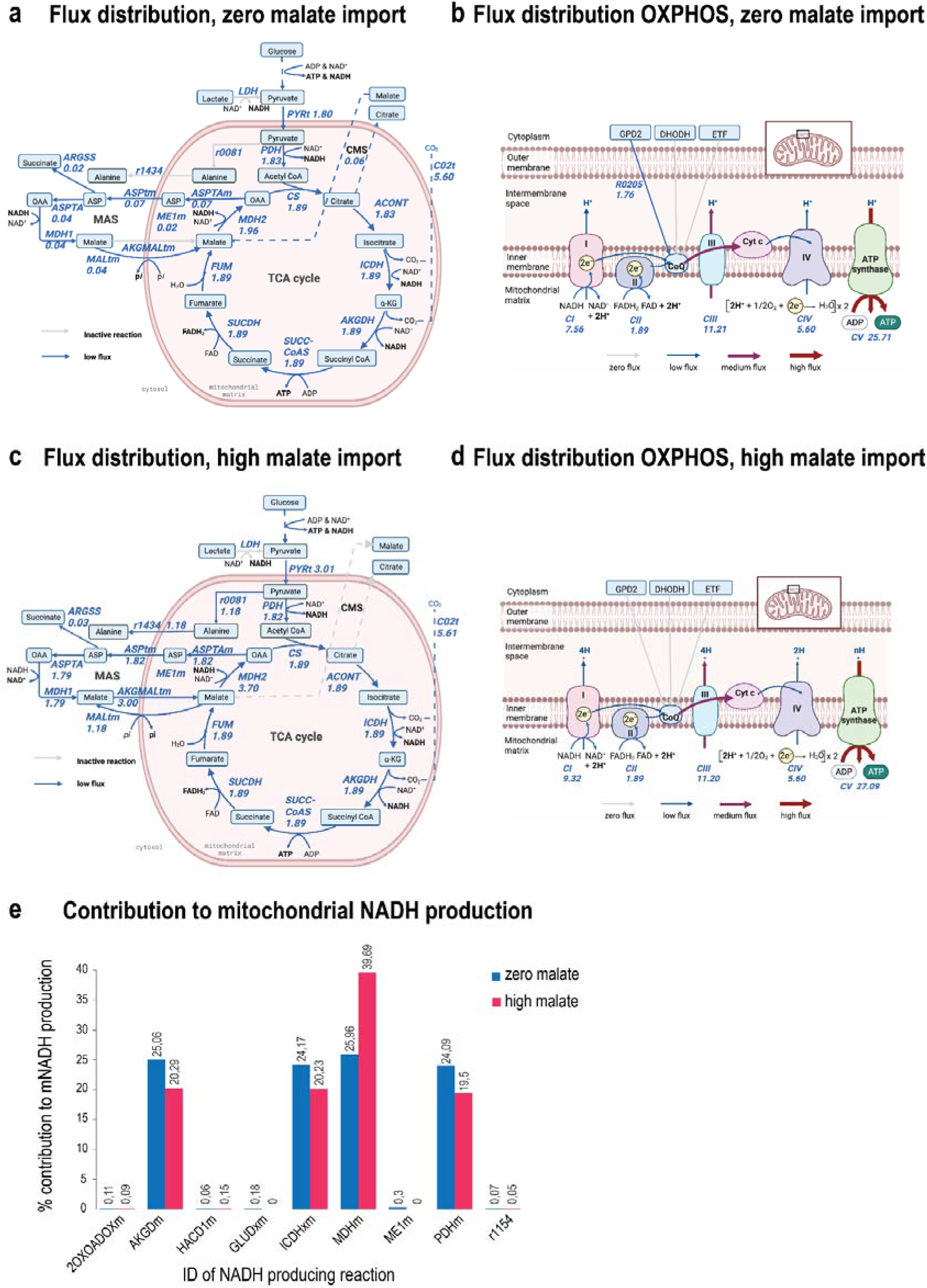
**(a)** Flux distribution through central carbon metabolism for blocked malate import (zero malate). Blocking malate import through MAS reroutes mitochondrial metabolism to induce glutamate import and activation of the citrate-malate shuttle which shuttles malate into the TCA cycle at the expense of citrate. Half of the importer malate is subsequently converted to pyruvate. Dashed lines represent multistep pathways carrying identical fluxes. **(b)** Flux distribution through OXPHOS for zero malate import. Blocking malate import through MAS reroutes is predicted to activate GPD2 reduction of CoQ complex in addition to the activity of Complex I (CI) and II (CII). This leads to the prediction of larger fluxes passing through CIII and CIV which along with CI generate a proton gradient and thus drive ATP synthesis from CV (ATP synthase). **(c)** Flux distribution through central carbon metabolism for high malate import. Flux distribution from allowing malate import at 3.00 (mmol/gDw/h). Dashed lines represent multiple metabolic reactions carrying identical fluxes. **(d)** Flux distribution through OXPHOS for high malate import. Malate import activates alternative CoQ-reducing pathways and higher ATP yields. (e) NADH production comparison between conditions.

We next analysed the predicted fluxes arising from the reactions belonging to OXPHOS (Figure 3 b). At zero malate import, reducing power in the form of NADH and FADH2 supplied from the TCA cycle were predicted to feed into their respective complexes (complex I (CI) and complex II (CII)) of the (RC) and both complexes were predicted to contribute to the reduction of CoQ. Additional electrons were predicted to be injected into the RC and reduce CoQ from the oxidation of glycerol 3-phosphate. Flux continued throughout the RC towards CIII and onwards to Cytochrome C (Cyt C) and CIV. CI, CIII and CIV are coupled to hydrogen ion pumping into the intermembrane space, with the reduction of CIII contributing to greater H^+^ pumping. The flux of hydrogen into the intermembrane space was then predicted to be consumed by ATP synthase (CV) to produce ATP which out of this subset of reactions carries the greatest flux of 25.71 mmol/gDw/h (Figure 2 a). In this condition, the model predicted an optimal flux of ATP synthesis to be 29.32 mmol/gDw/h, with the remaining ATP-producing reactions being the SUCC-CoAS from the TCA cycle within the mitochondria and the glycolytic reactions by phosphoglucokinase and pyruvate kinase from the cytoplasm.

#### High malate import

At high malate import, malate was completely converted to oxaloacetate (OAA) and NADH, the CI substrate. Unlike the zero malate condition, the conversion of imported malate to pyruvate through the mitochondrial malic enzyme (ME1m) reaction was inactive. Instead, the model predicted increased pyruvate import into mitochondria as a consequence of increased glycolytic flux. The predominant flux of pyruvate was incorporated into the TCA cycle via acetyl-CoA facilitated by pyruvate dehydrogenase (PDH) and the remaining flux was diverted into alanine metabolism and exported out of the mitochondria. Contributions made by citrate synthase (CS) in metabolising acetyl-COA and OAA into citrate then sustained steady-state fluxes of the TCA cycle at values of 1.89 (mmol/gDw/h) through the subsequent TCA cycle enzymes involving aconitase (ACONT), isocitrate dehydrogenase (ICDH), alpha-ketoglutarate dehydrogenase (AKGDH), succinate CoA synthetase (SUCC-CoAS), succinate dehydrogenase (SUCDH) and resulting in the production of the reductants NADH and succinate. The CMS was predicted to be inactive in this condition. The final TCA enzyme, fumarase (FUM) directed flux back to malate, and excess malate was then exported out of the mitochondria by MALtm, a malate/phosphate antiporter, resulting in the import of inorganic phosphate. Unconstraining the malate import reaction of the MAS led to higher residual fluxes being predicted from the MAS when being compared to blocking malate import and so greater fluxes were attributed to aspartate metabolism and export out of the mitochondria with subsequent conversion to oxaloacetate (OAA) via the GOT2 encoded reaction (ASPTA).

When analysing the flux through OXPHOS reactions at high malate, we found that forcing malate import through the MAS-mediated import reaction yielded increased CI reducing power within the mitochondria as a result of increased flux through the MDH2 reaction, yet yielded the same flux of the succinate oxidation through CII as predicted with the condition. As such, an increased flux was directed to complex I (CI), and flux through CII was the same in both conditions. Similarly, both CI and CII were predicted to reduce CoQ, and electron flux was passed along the RC to complex III (CIII), cytochrome C (Cyt C) and into complex IV (CIV). Involvement of GPD2 reduction of CoQ was this time predicted to be inactive. As a result of a greater flux through CI, its coupling with proton pumping and its contribution to the generation of the proton motive force, a greater flux through complex V (ATP synthase) was predicted (27.09 mmol/gDw/h). This, plus joint contributions of the other ATP-producing reactions (SUCCOASM, phosphoglucokinase and pyruvate kinase) led to a higher prediction of optimal ATP yields of 30.67 mmol/gDw/h.

The MAS is a well-known shuttle that increases levels of the CI reductant within the mitochondria. MitoMouse has predicted that inhibiting malate import through the MAS activates alternative metabolic pathways that produce NADH and so compensates for the reduced reductant production capacity. We next wanted to investigate all the other reactions that contribute to increasing mitochondrial NADH production, as a consequence of blocking or allowing malate import through the MAS. As seen in Figure 3 e, there are a total of 9 reactions that contributed to NADH production: 2-Oxoadipate:lipoamde 2-oxidoreductase (2OXOADOXm, involved in lysine metabolism), alpha-ketoglutarate dehydrogenase (AKGDm, TCA cycle), 3-Hydroxybutanoyl-CoA: NAD+ oxidoreductase (HACD1m, tryptophan metabolism), glutamate dehydrogenase (GLUDxm, glutamate metabolism), ICDHxm (Isocitrate dehydrogenase, TCA cycle), malate dehydrogenase (MDHm, TCA cycle), malic enzyme (ME1m, pyruvate metabolism), pyruvate dehydrogenase (PDHm, glycolysis/gluconeogenesis) and reaction 1154 (involved in threonine and methionine metabolism).

Upon inducing malate uptake through MAS, the only flux that strongly increased was through MDHm in the TCA cycle, which then contributed to nearly 40%, and therefore the majority of overall NADH production. The remaining 60% was similarly distributed and slightly reduced as compared to zero malate import (Figure 3 e). The slight increase of flux through all remaining reactions at zero malate import therefore seemed sufficient for NADH production and for compensating for the lower flux through the MDHm reaction (Figure 3 e).

### Flux variability analysis to determine essential reactions for each condition

FBA returns a single flux distribution that has satisfied the given constraints in maximising the objective function, but multiple flux distributions can exist that could still achieve the same objective. To this end, we performed Flux Variability Analysis (FVA) to characterise the entire range of feasible minimal and maximal fluxes through each of the reactions. A reaction essentiality analysis that combined the obtained fluxes resulting from FBA and FVA was performed to identify reactions as being either essential (E), substitutable (S) or blocked (B) when malate import was blocked, and when malate import was forced through the MAS. The aim of this was to further investigate how essential, or not, are the selected reactions from the predicted fluxes arising from FBA for the two respective conditions. The essentiality status of the top 20 reactions that exhibited the greatest absolute difference in flux between the two conditions are listed in Table 1.

**Table 1:** The top 20 scoring reactions, ordered by decreasing absolute difference in flux obtained when the model was constrained to inhibit malate import through the malate aspartate shuttle, and when malate import was constrained to its maximal value of 3 mmol/gDw/h. For each reaction, the flux resulting from FBA from each condition is presented along with the corresponding absolute difference in flux, and the essentiality status (E=essential, S=substitutable, B=blocked) of each reaction obtained from FVA.

| reaction | flux value<br>0<br>mmol/gDw/h<br>malate import | flux value<br>3<br>mmol/gDw/h<br>malate import | abs<br>difference | status for 0<br>mmol/gDw/h<br>malate import | status for 3<br>mmol/gDw/h<br>malate import |
| --- | --- | --- | --- | --- | --- |
| AKGMALtm | 0 | 3 | 3 | B | E |
| Hct_mitoMouse | 0.13 | 2.97 | 2.84 | E | E |
| Hmt_mitoMouse | -0.15 | -2.99 | 2.83 | E | E |
| Cl_mitoMouse | 7.56 | 9.32 | 1.76 | E | E |
| G3PD1 | -1.76 | 0.00 | 1.76 | E | S |
| r0205 | 1.76 | 0.00 | 1.76 | E | S |
| ASPGLUmB_mitoMouse | 0.07 | 1.82 | 1.75 | E | E |
| ASPTAm | -0.07 | -1.82 | 1.75 | E | E |
| ASPTA | 0.05 | 1.79 | 1.75 | E | E |
| MDH | -0.05 | -1.79 | 1.75 | E | E |
| MDHm | 1.96 | 3.70 | 1.74 | E | E |
| H2Otm | -33.13 | -34.52 | 1.39 | E | E |
| CVmitoMouse | 25.71 | 27.09 | 1.38 | E | E |
| ATPtmB_mitoM | 27.58 | 28.95 | 1.37 | S | S |
| ouse |  |  |  |  |  |
| EX_biomass_e | 29.32 | 30.67 | 1.36 | E | E |
| OF_ATP_mitoM<br>ouse | 29.32 | 30.67 | 1.36 | E | E |
| Biomass_mitoM<br>ouse | -29.32 | -30.67 | 1.36 | E | E |
| MALtm | 0.03 | -1.18 | 1.21 | S | S |
| PYRt2m | 1.80 | 3.01 | 1.21 | E | E |
| GLUt2mB_mito<br>Mouse | 0.00 | 1.19 | 1.19 | S | E |

For the zero-malate condition, out of the full complement of 560 reactions, 136 reactions were predicted to be essential, 124 reactions were blocked and the vast majority (∼53 %) of remaining reactions were substitutable. For the condition in which malate was forced through the MAS, 133 reactions were essential, 110 were blocked and the remaining 316 reactions were substitutable (Supplementary Table S1 c).

In both conditions, fluxes through the TCA cycle and flux sustained through all five complexes of the ETC were classified as essential. CI and CV were amongst the top 20 reactions to have the largest absolute difference in flux, whilst the fluxes obtained for the other complexes were similar in both conditions.

As observed in Table 1, reactions that exhibited different essentiality statuses between the two conditions are GPD2 (Glycerol-3-phosphate dehydrogenase), involved in phosphoglycerolipid metabolism which was predicted to be essential when malate import was blocked and otherwise substitutable, along with reaction R0205 which oxidises glycerol 3-phosphate and reduces CoQ. The only other reaction whose flux capacity was predicted to contribute the greatest to the differences observed between the conditions was the malate import reaction which was intentionally manipulated as a constraint. Citrate import into the TCA cycle via reaction R0205 was predicted to be active when malate import was blocked, and inactive when malate uptake was forced. In both cases, this reaction was predicted to be substitutable as the variability of flux for both conditions could be zero or non-zero.

Out of the subset of NADH-producing reactions, the majority of reactions were predicted to be essential in contributing to the switch of electron transport between CI and CII, as a result of malate import, however reactions GLUDxm and ME1m were both substitutable in both conditions.

## Discussion

We have here presented the most up-to-date and precise metabolic model for the mitochondrial metabolism of mouse cardiomyocytes, mitoMouse. Because of the similarity between mouse and human mitochondrial genomes, mitoMouse was built from the human mitoCore model by translating human genes to their mouse orthologs. It was recently discovered in intact mitochondria from murine mouse tissue that imported malate via MDH1 modulates the RC electron fuelling and triggers a switch from CII to CI reduction of CoQ, we refined the model by adding the newest information on CoQ reduction by the DHODH pathway; and we updated mitoMouse to SBML3. We have also provided a documented Jupyter notebook with detailed information on how to replicate the results presented in this work, making it therefore easily accessible and available without a licence to the research community. We tested mitoMouse for its accuracy in predicting realistic ATP yields and simulated the model using flux balance analysis to capture the switch of electron transfer to CoQ from CII to CI that arises from increased malate import into the TCA cycle via the Malate Aspartate Shuttle (MAS). Furthermore, we analysed the flux distributions resulting from zero to high malate import using Flux Variability Analysis. Finally, we identified the essentiality of reactions from the two conditions based on the minimum and maximal flux values obtained by FVA.

In the attempt to model and predict mitochondrial metabolism, organismal system-level models of metabolism that include mitochondria have been built, yet a species-specific model for mitochondrial metabolism has so far only been done for human^38^. As mouse is an important model organism that is often used to study mitochondrial bioenergetics and metabolism, this model represents an important building block to understand murine mitochondria. FBA on accurate metabolic models like mitoMouse is moreover helpful in deciphering potential underlying mechanisms under certain constrained conditions, and therefore can help in the design of novel experiments. Our model for instance predicted that by blocking malate import, compensatory reactions were activated such as the citrate-malate shuttle (Figure 3 a) and several NADH-producing pathways (Figure 3 e) to re-establish a balanced redox state. One could argue this could be a result of the optimisation procedure as the model responds to low fluxes of malate within the TCA cycle by increasing the import of malate from alternative sources. Adding confidence to this prediction is the fact that mitochondria that are experiencing an impaired redox state can disconnect fatty acid oxidation from the TCA cycle and instead, connect it to the citrate-malate shuttle^50^. The citrate-malate shuttle was also observed to be overexpressed in mitochondria of cancer cells that exhibit an imbalanced redox state^51^. The consensus from these studies is that the citrate-malate shuttle plays a crucial role in maintaining mitochondrial integrity by allowing the continual oxidation of fatty acids at times of redox imbalance, e.g. from inhibiting malate uptake. The substitutable nature of this shuttle, however, as predicted by FVA, highlights a non-essential role of the shuttle which could explain why there are only a few existing studies on this subject. This could be further investigated *in silico*, as well as experimentally.

In each of the tested conditions, metabolic fluxes were sustained through all five complexes involved in OXPHOS, whose reactions were classified as essential (Supplementary Table S1 c). This was because the capacity of each complex to carry flux was non-variable. These predictions are therefore overly simplistic for such a complex system that is heavily regulated by, and itself regulates metabolism. The essentiality of the CoQ complex^52^ and CV^53^ are well established in directing optimal electron flux to drive ATP synthesis. However, it was recently discovered that the complexes of the RC can arrange into different supercomplexes (SCs) depending on the availability of reductant. When NADH is available, CI forms a supercomplex with CIII and CIV; yet CII can form a supercomplex with CIII and IV when FADH_2_ is available^54^. Therefore, CI or CII can be dispensable for ATP production depending on reductant availability. Figure 3e shows the compensatory NADH-producing reactions that have gained in flux capacity as a result of inhibiting malate import. But both conditions result in flux being carried from the TCA cycle to generate FADH_2._ As a consequence, both substrates are available and steady-state fluxes obtained by linear programming then dictates their involvement with OXPHOS in order to satisfy the objective of optimised ATP production. The idea of modelling supercomplex assembly to more faithfully represent this switch of electron transfer to coQ is very interesting and novel and presents a future opportunity for mitoMouse.

The two reactions that exhibited different essentiality statuses between the two conditions are GPD2 (Glycerol-3-phosphate dehydrogenase) which produces glycerol-3-phosphate, and reaction R0205 which oxidises glycerol 3-phosphate whilst reducing CoQ (Table 1 and Figure 3 a). The activity of both these reactions was predicted to be essential to support ATP production when malate import was blocked and otherwise substitutable. GPD2 is an integral component of the RC and the glycerophosphate shuttle and so interconnects mitochondrial and cytosolic processes in mammals. Metabolic roles of GPD2 have been attributed to the re-oxidation of cytosolic NADH in glycolytic cells and to bypassing CI during cytosolic NADH oxidation^55^. By optimising ATP production, the model has indeed predicted the activation of GPD2 at times of high flux through CII and low flux through CI. This prediction is in agreement with ^55^, who report that electrons from glycerol-3-phosphate can entirely bypass CI. Nilsson et al.^56^ have also predicted the metabolic bypass of CI at times of optimal ATP production in mitochondria from muscle fibres. In their study, using a small-scale metabolic model of human muscle fibre metabolism, the authors predict a higher capacity of the glycerophosphate shuttle compared to the MAS for optimal ATP synthesis. The combined results of our prediction and published research reveals a seemingly important role associated with the activation of GPD2 in response to altered redox states and an effect in regulating the RC. This likewise could be explored further experimentally.

We want to stress that we here show only one possible application of mitoMouse, using one objective function (maximising ATP production) and applying it to a single experimental setup (increasing malate import). Many scenarios and experimental conditions can be modelled using mitoMouse in combination with FBA and FVA, by using different objective functions, or by interfering with different parts of the mitochondrial metabolic network (e.g. in mutant conditions by blocking certain parts of the metabolic network). Moreover, experimental data in the form of gene or protein expression, as well as different metabolic data can be used in the combination with mitoMouse, making the model generally very useful in predicting the effects of different cellular conditions on mitochondrial metabolism. MitoMouse is open source and freely available to the research community available here: https://gitlab.com/habermann_lab/mitomouse.

## DATA AVAILABILITY

mitoMouse is freely available at: https://gitlab.com/habermann_lab/mitomouse

## AUTHOR CONTRIBUTIONS

SPC and BHH conceived the study, SPC created the mitoMouse model with help from BHH. SPC was solely responsible for code development, FBA and FVA analysis. Results were interpreted by SPC, AM and BHH. The manuscript was written by SPC and BHH, with contributions from TM and AM.

## ACKNOWLEDGEMENTS

We would like to thank Jean-Marc Schwartz, Helene Puccio, the Habermann and Mourier teams for helpful feedback on this work.

## SUPPLEMENTARY MATERIALS

**Supplementary Table S1a :** metabolites and attributes of MitoMouse

**Supplementary Table S1b:** reactions and attributes of MitoMouse

**Supplementary Table S1c :** FBA, FVA and reaction essentiality analysis of low, and high malate import

**Supplementary File S1:** mitoMouse in XML format

**Supplementary File S2:** Jupyter notebook presenting and replicating results obtained in this work

