## Supplementary File S2 for "MitoMouse is a model reconstruction of murine mitochondrial metabolism": Chapman_etal_SupplFileS2_JupyterNotebook.ipynb

```
# %% [markdown]
**script to report fluxes for when malate import is constrained to 0mmol and 3mmol then FBA and FVA are performed and escher used to visualise flux distribtions**

# %% [markdown]
These simluaitons were performed using MacOS Bug SUr version 11.5.2, in conjunction with Python 3 and cobraPY using
the GLPK solver to perfrom FBA and FVA. Differences in the above may change the obtained results. The user will first need to install COBRApy. (https://github.com/opencobra/cobrapy/blob/devel/INSTALL.rst) and Escher (https://pypi.org/project/Escher/).

# %% [markdown]
To start, uplaod the model and this Jupyter notebook to the same location and make sure you know the path to the model.

# %% [code]
import cobra
import logging
import sys
import matplotlib.pyplot as plt
import escher
import numpy as np
import pandas as pd

# %% [code]
from os.path import join
from cobra.util import create_stoichiometric_matrix
from cobra.util.solver import linear_reaction_coefficients
from cobra.io import read_sbml_model, write_sbml_model
from cobra.flux_analysis import pfba
from escher import Builder
from time import sleep
from cobra.flux_analysis import flux_variability_analysis

# %% [code]
You now have to enter the path to the model within the "" of the following command.

# %% [code]
data_dir= "/home/stephen/Documents/research/FBA_mitochondria/FBA/arnaud_mourier_malate_prioritises_complex_1"
#make sure the directory to the where the model is saved on your comptuer is correct!!!

# %% [code]
model = cobra.io.read_sbml_model(join(data_dir, "mitoMouse_V3.xml")) #used for glucose and model exploration

# %% [code]
type(model.solver) #view the solver being implemented

# %% [code]
#explores the model and prints the number of metabolites, reactions, groups and comparetments present in th model
model

# %% [code]
print(len(model.genes))

# %% [code]
#reactuions and attributes can be ontained by parsing the "get_by_id() method"
model.reactions.get_by_id("PGI")

# %% [code]
#reactions and metabolites can be accessed by their index too
model.reactions[20]
model.metabolites[56]

# %% [code]
#printing exchange reactions
for b in model.boundary:
    print(b)

# %% [markdown]
## Performing FBA on the base model to predict ATP production with glucose metabolism (Figure 2b) #


# %% [markdown]
# Set model constraints

# %% [code]
# simuate aerobic growth: set all uptake reactions to 0, except glucose and oxygen
model.reactions.EX_ala_L_e.bounds = (0,0)
model.reactions.EX_arg_L_e.bounds = (0,0)
model.reactions.EX_asn_L_e.bounds = (0,0)
model.reactions.EX_asp_L_e.bounds = (0,0)
model.reactions.EX_bhb_e.bounds = (0,0)
model.reactions.EX_cys_L_e.bounds = (0,0)
model.reactions.EX_gly_e.bounds = (0,0)
model.reactions.EX_glyc_e.bounds = (0,0)
model.reactions.EX_h_e.bounds = (0,0)
model.reactions.EX_hco3_e.bounds = (0,0)
model.reactions.EX_hdca_e.bounds = (0,0)
model.reactions.EX_his_L_e.bounds = (0,0)
model.reactions.EX_ile_L_e.bounds = (0,0)
model.reactions.EX_lac_L_e.bounds = (0,0)
model.reactions.EX_leu_L_e.bounds = (0,0)
model.reactions.EX_lys_L_e.bounds = (0,0)
model.reactions.EX_pro_L_e.bounds = (0,0)
model.reactions.EX_ser_L_e.bounds = (0,0)
model.reactions.EX_so3_e.bounds = (0,0)
model.reactions.EX_thr_L_e.bounds = (0,0)
model.reactions.EX_val_L_e.bounds = (0,0)

model.reactions.EX_glc_D_e.bounds = (-100,0)
model.reactions.EX_o2_e.bounds = (-100,0)

# %% [markdown]
# Setting the objective function: the reaction which will be optimised #

# %% [code]
model.objective = model.reactions[70]
linear_reaction_coefficients(model) #reminds us of the objective function we have selected

# %% [markdown]
# Perform FBA on the model subject to impose constraints and optimsiing chosen objective function #

# %% [code]
realATP = model.optimize()
print(realATP) #prints the optial flux through the objective reaction

# %% [markdown]
# Plotting optimal ATP poroduction as a function of glucose uptake (Figure 2b) #

# %% [code]
glucose_transport = []#model.reaction[26]
ATP_production = []#model.reaction[70]


for i in np.arange(0,5,0.5): 
 
	model.reactions[26].lower_bound = - i
	model.reactions[26].upper_bound = i
	ATPsolution = model.optimize()         
	ATP_production.append(ATPsolution.objective_value)
	glucose_transport.append(model.reactions[26].lower_bound)

plt.plot(glucose_transport, ATP_production)

plt.ylabel('ATP production')
plt.xlabel('Glucose transport')
plt.show()

# %% [code]
#to improve clarty of the graph, we need to invert X axis, and remove the negative - Exchange reactions are modelled
#as negative uptake, so -30 means -30 uptake into mitochondria.

new_glc_transport = np.arange(0,5,0.5)

plt.plot(new_glc_transport, ATP_production, color='red')
plt.ylabel('ATP production (mmol gDw$^{-1}$ hr$^{-1}$)', fontsize=14)
plt.xlabel('Glucose transport into cytosol (mmol gDw$^{-1}$ hr$^{-1}$)', fontsize=14)
plt.show()

# %% [markdown]
## FBA to capture switch of COQ reduction as a function of malate import

# %% [code]
low_model = cobra.io.read_sbml_model(join(data_dir, "mitoMouse_V3.xml"))
high_model = cobra.io.read_sbml_model(join(data_dir, "mitoMouse_V3.xml"))

# %% [code]
#set all uptake reactions to 0, except glucose and oxygen, constrain malate import to 0 and also ETF fluxes
low_model.reactions.EX_acac_e.bounds = (0,0)
low_model.reactions.EX_ala_L_e.bounds = (0,0)
low_model.reactions.EX_arg_L_e.bounds = (0,0)
low_model.reactions.EX_asn_L_e.bounds = (0,0)
low_model.reactions.EX_asp_L_e.bounds = (0,0)
low_model.reactions.EX_bhb_e.bounds = (0,0)
low_model.reactions.EX_cys_L_e.bounds = (0,0)
low_model.reactions.EX_gly_e.bounds = (0,0)
low_model.reactions.EX_glyc_e.bounds = (0,0)
low_model.reactions.EX_h_e.bounds = (0,0)
low_model.reactions.EX_hco3_e.bounds = (0,0)
low_model.reactions.EX_hdca_e.bounds = (0,0)
low_model.reactions.EX_his_L_e.bounds = (0,0)
low_model.reactions.EX_ile_L_e.bounds = (0,0)
low_model.reactions.EX_lac_L_e.bounds = (0,0)
low_model.reactions.EX_leu_L_e.bounds = (0,0)
low_model.reactions.EX_lys_L_e.bounds = (0,0)
low_model.reactions.EX_pro_L_e.bounds = (0,0)
low_model.reactions.EX_ser_L_e.bounds = (0,0)
low_model.reactions.EX_so3_e.bounds = (0,0)
low_model.reactions.EX_thr_L_e.bounds = (0,0)
low_model.reactions.EX_val_L_e.bounds = (0,0)

#constrain all ETF fluxes to zero
low_model.reactions[143].bounds = (0,0)
low_model.reactions[149].bounds = (0,0)
low_model.reactions[154].bounds = (0,0)
low_model.reactions[159].bounds = (0,0)
low_model.reactions[164].bounds = (0,0)
low_model.reactions[169].bounds = (0,0)
low_model.reactions[212].bounds = (0,0)
low_model.reactions[227].bounds = (0,0)
low_model.reactions[263].bounds = (0,0)
low_model.reactions[282].bounds = (0,0)

#block uptake of malate through MAS
low_model.reactions.AKGMALtm.bounds = (0,0)

#allow glucse and oxygen uptake
low_model.reactions.EX_glc_D_e.bounds = (-1000,0)
low_model.reactions.EX_o2_e.bounds = (-1000,0)

# %% [code]
print ((low_model.reactions.EX_glc_D_e.bounds), "glucose import bounds")
print ((low_model.reactions.EX_o2_e.bounds), "oxygen import bounds")
print ((low_model.reactions.AKGMALtm.bounds), "Malate import reaction")

# %% [code]
low_model.objective = low_model.reactions[70]
linear_reaction_coefficients(low_model)

# %% [code]
low_malate = low_model.optimize()
print(low_malate) #returns flux through objective function

# %% [markdown]
## FBA to show that malate import triggers activation of complex II  and reduction of complex I flux (Figure 2c)

# %% [code]
malate_transport = []#model.reactions[131]
complex_1 = []#model.reactions[107]
complex_2 = []#model.reaction[108]

for i in np.arange(0,3.1,.1): #this increases the value of 'i' from 0 to 3, in 0.1 increments
    low_model.reactions[131].bounds = (i, i) #sets lower and upper bounds to the value of 'i'
    low_model.objective = low_model.reactions[70] #selects objective function
    solution = low_model.optimize()  #FBA optimisation       
    malate_transport.append(solution.fluxes[131]) #appends the flux through malate import reaction to a list 
    complex_1.append(solution.fluxes[107]) #same for flux through C1
    complex_2.append(solution.fluxes[108]) #same for flux through C2

# %% [code]
# create figure and axis objects with subplots()
fig,ax = plt.subplots() # we are plotting two plots on the same axis so need subplots
ax.plot(malate_transport,complex_1,color="#DF6589f7",marker="o")
ax.set_xlabel("malate import through MAS", fontsize = 14)
ax.set_ylabel("complex 1 flux",color="#DF6589f7",fontsize=14)

# %% [code]
# twin object for two different y-axis on the sample plot
ax2=ax.twinx()
# make a plot with different y-axis using second axis object
ax2.plot(malate_transport,complex_2,color="#3C1053FF",marker="o")
ax2.set_ylabel("complex 2 flux",color="#3C1053FF",fontsize=14)
fig.show()
# save the plot as a file
fig.savefig('malate_rewires_electron_transport.jpg',format='jpg', dpi=500) 
#the figure is saved as jpg in the same folder that contains this jupyter notebook

# %% [code]
print(complex_1)
print('#######################')
print(complex_2)
print('#######################')
print(malate_transport)

# %% [code]
print(low_malate[107], model.reactions[107].id, model.reactions[107].reaction)
print(low_malate[108], model.reactions[108].id, model.reactions[108].reaction)
print(low_malate.fluxes[109], model.reactions[109].id, model.reactions[109].reaction)
print(low_malate.fluxes[110], model.reactions[110].id, model.reactions[110].reaction)
print(low_malate.fluxes[111], model.reactions[110].id, model.reactions[110].reaction)

# %% [code]
#rest all constraints as the previsu simualtion updated the consraints - for comparison
low_model.reactions.EX_acac_e.bounds = (0,0)
low_model.reactions.EX_ala_L_e.bounds = (0,0)
low_model.reactions.EX_arg_L_e.bounds = (0,0)
low_model.reactions.EX_asn_L_e.bounds = (0,0)
low_model.reactions.EX_asp_L_e.bounds = (0,0)
low_model.reactions.EX_bhb_e.bounds = (0,0)
low_model.reactions.EX_cys_L_e.bounds = (0,0)
low_model.reactions.EX_gly_e.bounds = (0,0)
low_model.reactions.EX_glyc_e.bounds = (0,0)
low_model.reactions.EX_h_e.bounds = (0,0)
low_model.reactions.EX_hco3_e.bounds = (0,0)
low_model.reactions.EX_hdca_e.bounds = (0,0)
low_model.reactions.EX_his_L_e.bounds = (0,0)
low_model.reactions.EX_ile_L_e.bounds = (0,0)
low_model.reactions.EX_lac_L_e.bounds = (0,0)
low_model.reactions.EX_leu_L_e.bounds = (0,0)
low_model.reactions.EX_lys_L_e.bounds = (0,0)
low_model.reactions.EX_pro_L_e.bounds = (0,0)
low_model.reactions.EX_ser_L_e.bounds = (0,0)
low_model.reactions.EX_so3_e.bounds = (0,0)
low_model.reactions.EX_thr_L_e.bounds = (0,0)
low_model.reactions.EX_val_L_e.bounds = (0,0)

#constrain all ETF fluxes to zero
low_model.reactions[143].bounds = (0,0)
low_model.reactions[149].bounds = (0,0)
low_model.reactions[154].bounds = (0,0)
low_model.reactions[159].bounds = (0,0)
low_model.reactions[164].bounds = (0,0)
low_model.reactions[169].bounds = (0,0)
low_model.reactions[212].bounds = (0,0)
low_model.reactions[227].bounds = (0,0)
low_model.reactions[263].bounds = (0,0)
low_model.reactions[282].bounds = (0,0)

#block uptake of malate through MAS
low_model.reactions.AKGMALtm.bounds = (0,0)

#allow glucse and oxygen uptake
low_model.reactions.EX_glc_D_e.bounds = (-1000,0)
low_model.reactions.EX_o2_e.bounds = (-1000,0)

# %% [code]
low_model.objective = low_model.reactions[70]
linear_reaction_coefficients(low_model)

# %% [code]
low_malate = low_model.optimize()
print(low_malate) #returns flux through objective function

# %% [markdown]
### Constraints for when malate import through MAS is constrained to 3mmol /gDw/hr for comparison with low (blocked) malate import

# %% [code]
#set all uptake reactions to 0, except glucose and oxygen, constrain malate import to 0 and also ETF fluxes
high_model.reactions.EX_acac_e.bounds = (0,0)
high_model.reactions.EX_ala_L_e.bounds = (0,0)
high_model.reactions.EX_arg_L_e.bounds = (0,0)
high_model.reactions.EX_asn_L_e.bounds = (0,0)
high_model.reactions.EX_asp_L_e.bounds = (0,0)
high_model.reactions.EX_bhb_e.bounds = (0,0)
high_model.reactions.EX_cys_L_e.bounds = (0,0)
high_model.reactions.EX_gly_e.bounds = (0,0)
high_model.reactions.EX_glyc_e.bounds = (0,0)
high_model.reactions.EX_h_e.bounds = (0,0)
high_model.reactions.EX_hco3_e.bounds = (0,0)
high_model.reactions.EX_hdca_e.bounds = (0,0)
high_model.reactions.EX_his_L_e.bounds = (0,0)
high_model.reactions.EX_ile_L_e.bounds = (0,0)
high_model.reactions.EX_lac_L_e.bounds = (0,0)
high_model.reactions.EX_leu_L_e.bounds = (0,0)
high_model.reactions.EX_lys_L_e.bounds = (0,0)
high_model.reactions.EX_pro_L_e.bounds = (0,0)
high_model.reactions.EX_ser_L_e.bounds = (0,0)
high_model.reactions.EX_so3_e.bounds = (0,0)
high_model.reactions.EX_thr_L_e.bounds = (0,0)
high_model.reactions.EX_val_L_e.bounds = (0,0)

#constrain all ETF fluxes to zero
high_model.reactions[143].bounds = (0,0)
high_model.reactions[149].bounds = (0,0)
high_model.reactions[154].bounds = (0,0)
high_model.reactions[159].bounds = (0,0)
high_model.reactions[164].bounds = (0,0)
high_model.reactions[169].bounds = (0,0)
high_model.reactions[212].bounds = (0,0)
high_model.reactions[227].bounds = (0,0)
high_model.reactions[263].bounds = (0,0)
high_model.reactions[282].bounds = (0,0)

#constrain uptake of malate through MAS
high_model.reactions.AKGMALtm.bounds = (3,3)

#allow glucose and oxygen uptake
high_model.reactions.EX_glc_D_e.bounds = (-1000,0)
high_model.reactions.EX_o2_e.bounds = (-1000,0)

# %% [code]
#check reactions bounnds have been implemented
print (high_model.reactions.EX_glc_D_e.bounds)
print (high_model.reactions.EX_o2_e.bounds)
print (high_model.reactions.AKGMALtm.bounds)


# %% [code]
high_model.objective = high_model.reactions[70]
linear_reaction_coefficients(high_model)

# %% [code]
high_malate = model.optimize()
print(high_malate)

# %% [markdown]
## Compare flux distributions obtained from FBA of low vs high malate import ##

# %% [code]
#MAS import: low
print(low_malate.fluxes[129], model.reactions[129].id, model.reactions[129].reaction)
print(low_malate.fluxes[130], model.reactions[130].id, model.reactions[130].reaction)
print(low_malate.fluxes[131], model.reactions[131].id, model.reactions[131].reaction)
print(low_malate.fluxes[132], model.reactions[132].id, model.reactions[132].reaction)
print(low_malate.fluxes[133], model.reactions[133].id, model.reactions[133].reaction)

# %% [code]
#MAS import: high
print(high_malate.fluxes[129], model.reactions[129].id, model.reactions[129].reaction)
print(high_malate.fluxes[130], model.reactions[130].id, model.reactions[130].reaction)
print(high_malate.fluxes[131], model.reactions[131].id, model.reactions[131].reaction)
print(high_malate.fluxes[132], model.reactions[132].id, model.reactions[132].reaction)
print(high_malate.fluxes[133], model.reactions[133].id, model.reactions[133].reaction)

# %% [code]
#OXPHOS fluxes: low
print(low_malate.fluxes[107], model.reactions[107].id, model.reactions[107].reaction)
print(low_malate.fluxes[108], model.reactions[108].id, model.reactions[108].reaction)
print(low_malate.fluxes[109], model.reactions[109].id, model.reactions[109].reaction)
print(low_malate.fluxes[110], model.reactions[110].id, model.reactions[110].reaction)
print(low_malate.fluxes[111], model.reactions[111].id, model.reactions[111].reaction)

# %% [code]
#OXPHOS fluxes: high
print(high_malate.fluxes[107], model.reactions[107].id, model.reactions[107].reaction)
print(high_malate.fluxes[108], model.reactions[108].id, model.reactions[108].reaction)
print(high_malate.fluxes[109], model.reactions[109].id, model.reactions[109].reaction)
print(high_malate.fluxes[110], model.reactions[110].id, model.reactions[110].reaction)
print(high_malate.fluxes[111], model.reactions[111].id, model.reactions[111].reaction)

# %% [code]
#priting all resulting fluxes for each reaction
print(low_malate.fluxes)

# %% [code]
print(high_malate.fluxes)

# %% [markdown]
## comparison of low vs high for reactions that contribute to NADHm production (Figure 2e)

# %% [code]
low_model.metabolites.nadh_m.summary()

# %% [code]
high_model.metabolites.nadh_m.summary()

# %% [code]
# data to plot all NADH producing reactions
n_groups = 9
low_malate = [0.11,25.06,0.06,0.18,24.17,25.96,0.30,24.09,0.07]
high_malate = [0.09,20.29,0.15,0.0,20.23,39.69,0.0,19.50,0.05]

# create plot
fig, ax = plt.subplots()
index = np.arange(n_groups)
bar_width = 0.35
opacity = 0.8

rects1 = plt.bar(index, low_malate, bar_width,
alpha=opacity,
color='b',
label='0 mmol malate')

rects2 = plt.bar(index + bar_width, high_malate, bar_width,
alpha=opacity,
color='g',
label='3mmol malate')

plt.xlabel('reaction ID')
plt.ylabel('contribution to NADH production (%)')
plt.title('NADH production')
plt.xticks(index + bar_width, ('2OXOADOXm', 'AKGDm', 'HACD1m', 'GLUDxm', 'ICDHxm', 'MDHm', 'ME1m', 'PDHm', 'r1154'), rotation='vertical')
plt.legend()

plt.tight_layout()
plt.show() #shows figure 2e

# %% [markdown]
## FVA for Reaction Essentiality Analysis for low malate import


# %% [code]
from cobra.flux_analysis import flux_variability_analysis

# %% [code]
#reload model
model= cobra.io.read_sbml_model(join(data_dir, "mitoMouse_V3.xml"))

# %% [code]
#apply constrains to et malate import to 0
model.reactions.EX_acac_e.bounds = (0,0)
model.reactions.EX_ala_L_e.bounds = (0,0)
model.reactions.EX_arg_L_e.bounds = (0,0)
model.reactions.EX_asn_L_e.bounds = (0,0)
model.reactions.EX_asp_L_e.bounds = (0,0)
model.reactions.EX_bhb_e.bounds = (0,0)
model.reactions.EX_cys_L_e.bounds = (0,0)
model.reactions.EX_gly_e.bounds = (0,0)
model.reactions.EX_glyc_e.bounds = (0,0)
model.reactions.EX_h_e.bounds = (0,0)
model.reactions.EX_hco3_e.bounds = (0,0)
model.reactions.EX_hdca_e.bounds = (0,0)
model.reactions.EX_his_L_e.bounds = (0,0)
model.reactions.EX_ile_L_e.bounds = (0,0)
model.reactions.EX_lac_L_e.bounds = (0,0)
model.reactions.EX_leu_L_e.bounds = (0,0)
model.reactions.EX_lys_L_e.bounds = (0,0)
model.reactions.EX_pro_L_e.bounds = (0,0)
model.reactions.EX_ser_L_e.bounds = (0,0)
model.reactions.EX_so3_e.bounds = (0,0)
model.reactions.EX_thr_L_e.bounds = (0,0)
model.reactions.EX_val_L_e.bounds = (0,0)

#constrain all ETF fluxes to zero
model.reactions[143].bounds = (0,0)
model.reactions[149].bounds = (0,0)
model.reactions[154].bounds = (0,0)
model.reactions[159].bounds = (0,0)
model.reactions[164].bounds = (0,0)
model.reactions[169].bounds = (0,0)
model.reactions[212].bounds = (0,0)
model.reactions[227].bounds = (0,0)
model.reactions[263].bounds = (0,0)
model.reactions[282].bounds = (0,0)


#block uptake of malate through MAS
model.reactions.AKGMALtm.bounds = (0,0)
model.reactions.EX_glc_D_e.bounds = (-1000,0)
model.reactions.EX_o2_e.bounds = (-1000,0)

# %% [code]
#set objective function
model.objective = model.reactions[70]
linear_reaction_coefficients(model)

# %% [code]
low_malate = model.optimize()
print(low_malate)

# %% [code]
low_malate_fva = flux_variability_analysis(model, model.reactions)

# %% [code]
print(low_malate_fva)

# %% [code]
for i in range(len(low_malate_fva)): 
    #print(low_malate_fva.maximum[i]) #prints the maximum flux carried by the reaction
    print(low_malate_fva.minimum[i]) #prints the minimum flux carried by the reaction

# %% [markdown]
# FBA for 3mmol malate import

# %% [code]
high_model

# %% [code]
#apply constrains to et malate import to 3
high_model.reactions.EX_acac_e.bounds = (0,0)
high_model.reactions.EX_ala_L_e.bounds = (0,0)
high_model.reactions.EX_arg_L_e.bounds = (0,0)
high_model.reactions.EX_asn_L_e.bounds = (0,0)
high_model.reactions.EX_asp_L_e.bounds = (0,0)
high_model.reactions.EX_bhb_e.bounds = (0,0)
high_model.reactions.EX_cys_L_e.bounds = (0,0)
high_model.reactions.EX_gly_e.bounds = (0,0)
high_model.reactions.EX_glyc_e.bounds = (0,0)
high_model.reactions.EX_h_e.bounds = (0,0)
high_model.reactions.EX_hco3_e.bounds = (0,0)
high_model.reactions.EX_hdca_e.bounds = (0,0)
high_model.reactions.EX_his_L_e.bounds = (0,0)
high_model.reactions.EX_ile_L_e.bounds = (0,0)
high_model.reactions.EX_lac_L_e.bounds = (0,0)
high_model.reactions.EX_leu_L_e.bounds = (0,0)
high_model.reactions.EX_lys_L_e.bounds = (0,0)
high_model.reactions.EX_pro_L_e.bounds = (0,0)
high_model.reactions.EX_ser_L_e.bounds = (0,0)
high_model.reactions.EX_so3_e.bounds = (0,0)
high_model.reactions.EX_thr_L_e.bounds = (0,0)
high_model.reactions.EX_val_L_e.bounds = (0,0)

#constrain all ETF fluxes to zero
high_model.reactions[143].bounds = (0,0)
high_model.reactions[149].bounds = (0,0)
high_model.reactions[154].bounds = (0,0)
high_model.reactions[159].bounds = (0,0)
high_model.reactions[164].bounds = (0,0)
high_model.reactions[169].bounds = (0,0)
high_model.reactions[212].bounds = (0,0)
high_model.reactions[227].bounds = (0,0)
high_model.reactions[263].bounds = (0,0)
high_model.reactions[282].bounds = (0,0)

#allow uptake of malate through MAS, constrain lower and upper bounds to 3
high_model.reactions.AKGMALtm.bounds = (3,3)

#allow uncosntrained glucose and oxygen uptake
high_model.reactions.EX_glc_D_e.bounds = (-1000,0)
high_model.reactions.EX_o2_e.bounds = (-1000,0)

# %% [code]
high_model.objective = high_model.reactions[70]
linear_reaction_coefficients(high_model)

# %% [code]
high_malate = model.optimize()
print(high_malate)

# %% [code]
high_malate_fva = flux_variability_analysis(model, model.reactions)

# %% [code]
print(high_malate_fva)

# %% [code]
for i in range(len(high_malate_fva)):
    print(high_malate_fva.minimum[i])
    #print(high_malate_fva.maximum[i])

# %% [markdown]
## Flux visulisation using ESCHER ##

# %% [code]
#Visualise fluxes
#pip install escher
import escher
from escher import Builder
import cobra
from time import sleep

builder = Builder() 
builder

# %% [code]
escher.list_available_models() #returns the names of available models 

# %% [code]
escher.list_available_maps() #returns the names of available metabolic maps

# %% [code]
#loading metabolic maps and models from Escher
builder = Builder(
    map_name='RECON1.Carbohydrate metabolism',
    model_name='RECON1'
)

builder 

# %% [markdown]
#escher visulasation of fluxes
builder.reaction_data = low_malate.fluxes
builder

# %% [code]
builder.reaction_data = high_malate.fluxes
builder

# %% [markdown]
note that here the  two maps show similar fluxes, and that is becuase the reported fluxes in Chapman et al. 2023 are simiar, and so requires the user to manually edit the visuation of the maps accordingly to show differential flux distribution. 

# %% [code]
```
